# A chromatin extension model for insulator function based on comparison of high-resolution chromatin conformation capture and polytene banding maps

**DOI:** 10.1101/129577

**Authors:** Michael R. Stadler, Michael B. Eisen

## Abstract

Insulator proteins bind to specific genomic loci and have been shown to play a role in partitioning genomes into independent domains of gene expression and chromatin structure. Despite decades of study, the mechanism by which insulators establish these domains remains elusive. Here, we use genome-wide chromatin conformation capture (Hi-C) to generate a high-resolution map of spatial interactions of chromatin from *Drosophila melanogaster* embryos. We show that from the earliest stages of development the genome is divided into distinct topologically associated domains (TADs), that we can map the boundaries between TADs to sub-kilobase resolution, and that these boundaries correspond to 500-2000 bp insulator elements. Comparing this map with a detailed assessment of the banding pattern of a region of a polytene chromosome, we show that these insulator elements correspond to low density polytene interbands that divide compacted bands, which correspond to TADs. It has been previously shown that polytene interbands have low packing ratios allowing the conversion of small genomic distances (in base pairs) into a large physical distances. We therefore suggest a simple mechanism for insulator function whereby insulators increase the physical space between adjacent domains via the unpacking and extension of intervening chromatin. This model provides an intuitive explanation for known features of insulators, including the ability to block enhancer-promoter interactions, limit the spread of heterochromatin, and organize the structural features of interphase chromosomes.

## Introduction

Beginning in the late 19th century, cytological investigations of the polytene chromosomes of insect salivary glands implicated the physical structure of interphase chromosomes in their metabolic functions (1–5). Progressively more detailed optical and electron microscopic analysis of polytene chromosomes in *Drosophila melanogaster* have identified a stereotyped banding pattern with compacted, DNA-rich “bands” alternating with extended, DNA-poor “interband” regions(6–10). While much is now known about the molecular properties of these two types of chromatin (11–20), the precise molecular nature of the banding structure remains elusive.

Several methods have been developed in the past decade for the isolation and high-throughput characterization of chromosomal regions that are colocalized within nuclei (21–24), yielding genome-wide maps of chromatin structure in a number of organisms and tissues. These experiments revealed that the interphase chromatin fiber is organized into topologically-associated domains (TADs), contiguous regions of the genome that exhibit enriched three-dimensional contacts between loci within the TAD, and depleted contacts between loci in different TADs. Eagen et al. recently showed that TADs identified from genome-wide chromatin conformation capture (Hi-C) at approximately 15 kb resolution from polytene nuclei of *Drosophila melanogaster* largely correspond to the bands of polytene chromosomes, and suggested that inter-TADs are regions of decompacted chromatin containing the promoters and regulatory elements of housekeeping genes (25).

We have been interested in the establishment of interphase chromatin structure during early *Drosophila* development because of its relationship to transcriptional activation (26–28), and have used deep sequencing of Hi-C libraries prepared with a 4-cutter restriction enzyme to generate a map of chromosome structure in cellular blastoderm embryos at sub-kilobase resolution. These data have many interesting features that we will present elsewhere, some of which have recently been discussed by (29,30). Here we focus exclusively on the relationship between our high-resolution Hi-C data, high-resolution polytene band patterns described by Vatolina et al. (31), and additional genomic data from our lab (26) and others (32–35) that shed light the molecular origins of chromosome organization.

## Results and Discussion

We performed *in situ* Hi-C (22) in the early *Drosophila melanogaster* embryo using the MboI restriction enzyme (^GATC) and sequenced ∼276 million productive linkages, a combination that yielded sub-kilobase resolution of chromatin architecture. From these data, we used methods similar to those employed by other groups (21,22,29,36,37) to identify regions of self-association (TADs) as well as boundary elements between those regions. Crucially, our data allowed the identification of TADs as small as a few kilobases. The boundary elements we identified between these TADs appear to correspond to classical insulator elements: regions of 500-2000 bp that are sensitive to DNAse digestion and are strongly bound by known insulator proteins. Noting that interband regions are also characterized by DNase hypersensitivity and insulator protein association, we wondered about the detailed relationship between our Hi-C TADs and boundaries and the banding structure observed in polytene chromosomes.

There is surprisingly little data associating features of polytene structure to the genome at high resolution. Vatolina et al. used exquisitely careful electron microscopy to identify the fine banding pattern of the 65 kb region between polytene bands 10A1-2 and 10B1-2, revealing that this region, which appears as a single interband under a light microscope, actually contains six faint bands and seven interbands (31). Vatolina et al. then used available molecular genomic data to infer genomic locations for these bands and interbands, and validated four of their interband assignments by FISH. We compared these inferred mappings of polytene banding patterns to our Hi-C data.

Figure 1 shows the correspondence between Vatolina et al.’s polytene map from this region and our high-resolution Hi-C data, along with measures of early embryonic DNase hypersentivity from (26) and the binding of four insulator proteins (32). Strikingly, there is a near-perfect correspondence between the assignments of Vatolina et al. and our Hi-C data: bands correspond to TADs, and interbands correspond to the insulator elements that separate the TADs. Eagen et al. previously identified a broad correspondence between polytene interbands and inter-TAD regions from their Hi-C data (25). However, our higher resolution Hi-C data reveals fine structure within these inter-TAD regions, and as a result we can now see a precise correspondence between insulators (regions of 500-2000 bp that are bound by insulator proteins and divide topological domains) and polytene interbands that has not, to our knowledge, been described previously.

**Figure 1:**
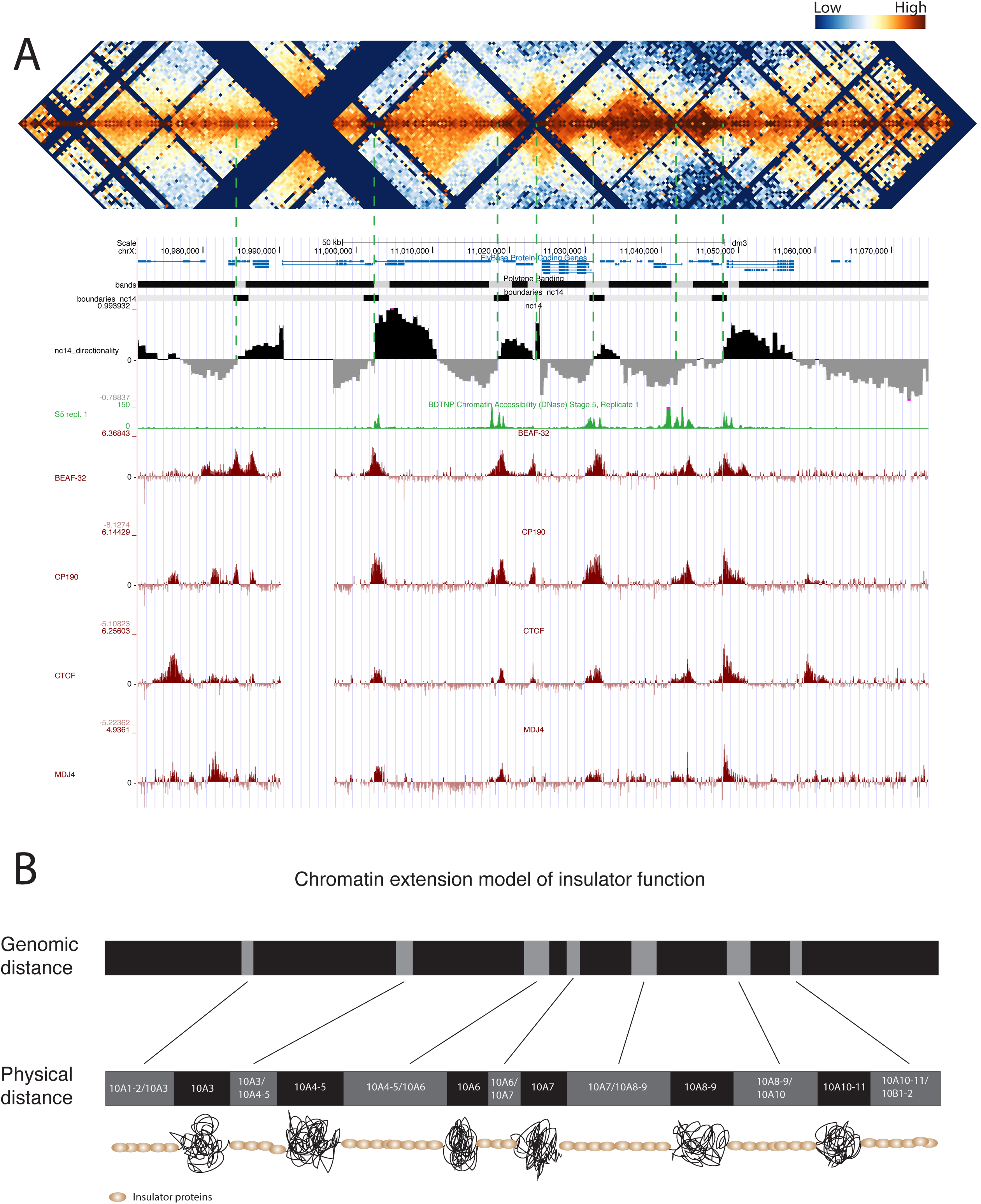
(A) Combined view from the UCSC genome browser of polytene banding structure resolved by (31), Hi-C interaction heatmap, Hi-C directionality index, DNase accessibility, ChIP-chip data for the insulator proteins CP190, BEAF-32, dCTCF, and Mod(mdg4) for the region covering chrX:10,971,551-11,074,797. Dashed green lines indicate the locations of TAD boundaries identified by Hi-C. Hi-C blocks lacking data result from genomic regions that lack MboI cut sites (GATC) and are uninformative. (B) Graphical representation of the chromatin expansion model of insulator function. The bands shown are proportionally sized according to genomic size (bp, top) and physical distance in polytene chromosomes (bottom) as measured by (31). Insulator proteins are represented to indicate that they drive the extension of these regions, not to imply that they coat them entirely. The presence of active promoters within the borders of a subset of these interbands suggests that these regions are not entirely occluded by insulator proteins. The mechanism(s) by which insulator proteins decompact chromatin are unclear.

A key feature that distinguishes polytene interbands from bands is their low compaction ratio: they span a larger physical distance per base pair. Our observation that insulators correspond so precisely to interbands suggest a simple and intuitive mechanism for insulator activity whereby insulators function by increasing the physical space between adjacent domains via the unpacking and extension of intervening chromatin.

Insulators have been associated with various functional properties. Principal among these, in addition to their role in organizing chromatin into domains, insulators block interactions between enhancers and promoters exclusively when inserted between them (38–44) and protect transgenes from position effect variegation and block the spread of chromatin silencing states (45–50). Without invoking complex interactions or chromosomal geometries, this chromatin extension model for insulator function can explain these defining characteristics via simple physical separation.

## Materials and Methods

Oregon R strain embryos were collected, aged, fixed with 5% PFA for 28 minutes, and hand sorted to isolate early cellularized (mid stage 5) embryos. In situ Hi-C was performed on 75-500 embryos as described in Rao et al. (22) with minor modifications, using the restriction enzyme MboI. Paired-end sequencing with 100 bp per side was performed on the Illumina Hi-Seq 2500. We prepared and sequenced libraries from seven independent biological and technical replicates. Combined data for all stage 5 samples is shown in this paper.

Sequenced reads were aligned to the flybase 5_22 genome (dm3) using bowtie (51), retaining only read pairs in which both ends mapped uniquely to the genome. Resulting reads were assessed and filtered for quality similar to (22). Linkage matrices at various resolutions were then generated by binning the genome into bins of different sizes. Local directionality scores for high-resolution (500 bp) bins were generated by comparing the number of upstream and downstream linkages within a 15 kb window for each bin. Boundary elements were identified heuristically by searching for sites of transitions between upstream-bias and downstream-bias and applying thresholds to identify boundaries with varying stringency.

ChIP data for insulator proteins was downloaded from the modEncode server and is described in (32,33,35). DNase-seq data was available at the UCSC genome browser (52,53)and was generated by (26).

